# A substrate-interacting region of Parkin directs ubiquitination of the mitochondrial GTPase Miro1

**DOI:** 10.1101/2024.06.03.597144

**Authors:** Joanna Koszela, Anne Rintala-Dempsey, Giulia Salzano, Viveka Pimenta, Outi Kamarainen, Mads Gabrielsen, Aasna L. Parui, Gary S. Shaw, Helen Walden

## Abstract

Mutations in the gene encoding for the E3 ubiquitin ligase Parkin have been linked to early-onset Parkinson’s disease. Besides many other cellular roles, Parkin is involved in clearance of damaged mitochondria via mitophagy - a process of particular importance in dopaminergic neurons. Upon mitochondrial damage, Parkin accumulates at the outer mitochondrial membrane and is activated, leading to ubiquitination of many mitochondrial substrates and recruitment of mitophagy effectors. While the activation mechanisms of autoinhibited Parkin have been extensively studied, it remains unknown how Parkin recognises its substrates for ubiquitination, and no substrate interaction site in Parkin has been reported. Here, we identify a conserved region in the flexible linker between the Ubl and RING0 domains of Parkin, which is indispensable for Parkin interaction with the mitochondrial GTPase Miro1. Our results explain the preferential targeting and ubiquitination of Miro1 by Parkin and provide a biochemical explanation for the presence of Parkin at the mitochondrial membrane prior to activation induced by mitochondrial damage. Our findings are important for understanding mitochondrial homeostasis and may inspire new therapeutic avenues for Parkinson’s disease.

## Introduction

Heritable forms of Parkinson’s disease (PD) account for 5-10% of all PD cases. The implication of mutations in *PRKN* gene, encoding for the E3 ubiquitin ligase Parkin (Shimura *et al*, 2000; Zhang *et al*, 2000), in early-onset, autosomal recessive parkinsonism was described in 1998 (Kitada *et al*, 1998). Since then, intensive research efforts have been made to understand Parkin function and structure. With an increasing number of identified PD patient mutations throughout the *PRKN* sequence (Zhang *et al*, 2021; Li *et al*, 2021; Jiang *et al*, 2020; Kasten *et al*, 2018; Taghavi *et al*, 2018; Tan *et al*, 2019), it became clear that all domains, and even supposedly disordered regions of Parkin are important for its function. Parkin is a RING-BetweenRING-RING (RBR) E3 ubiquitin ligase (Wenzel *et al*, 2011), composed of five domains (Riley *et al*, 2013): an N-terminal ubiquitin-like domain (Ubl), connected by a 65-residue flexible region (linker) to a unique to Parkin RING0 domain, which, together with RING1 and BRcat (IBR) constitute a more rigid ‘core’ of Parkin (Fig. 1A). At the C-terminus, another partially disordered ‘tether’ region connects the ‘core’ to the catalytic Rcat domain of Parkin, containing the active site. Several elements in Parkin maintain its closed, autoinhibited conformation: Ubl blocks the predicted donor ubiquitin binding site, the Repressive Element of Parkin (REP) in the tether region blocks the incoming E2 enzyme binding site and RING0 partially occludes the active site cysteine in Rcat (Kumar *et al*, 2017, 2015; Chaugule *et al*, 2011; Trempe *et al*, 2013; Riley *et al*, 2013; Wauer *et al*, 2015). Parkin, like other E3s, catalyses the last step in the ubiquitination reaction and modifies target proteins on lysine residues with a small protein, ubiquitin. To achieve this, Parkin, as an RBR ligase, is thought to interact with an E2 enzyme charged with ubiquitin, accepts ubiquitin onto its active site, interacts with or position itself in sufficient proximity to a substrate and finally transfers ubiquitin onto a target lysine. To fulfil its role, Parkin must first be released from its autoinhibition.

**Figure 1:**
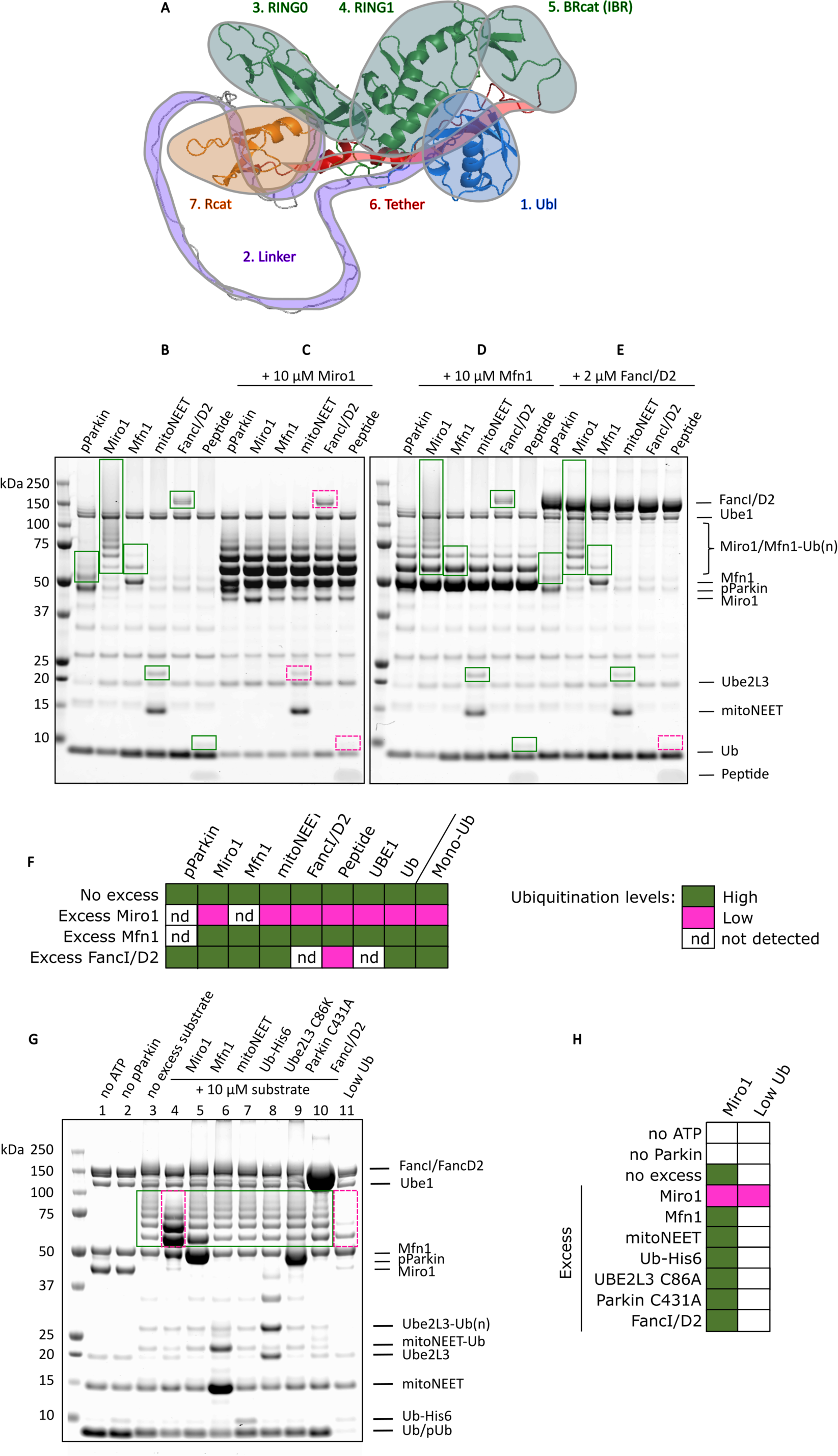
Active Parkin ubiquitinates various proteins but exhibits a preference towards Miro1 in a mix of substrates. A Schematic of Parkin in its autoinhibited conformation as predicted by ColabFold, with outlines of the structural elements in sequence order: Ubl (blue), linker (purple), RING0, RING1, BRcat (IBR) (green), tether (red), Rcat (orange). The prediction allows to appreciate the length of the linker as compared to Parkin domains. B: Ubiquitination assays were performed with pParkin and various proteins as substrates as indicated and resolved on SDS-PAGE. Represented are Coomassie-stained gels. Effects on substrate ubiquitination by pParkin were tested in excess of Miro1 (C), Mfn1 (D) or FancI/D2 (E). Green boxes indicate unaffected ubiquitination pattern, dashed pink boxes indicate decreased ubiquitination. F: Summary of results from B-E. G: Miro1 ubiquitination assay was performed with excess of various other substrates as indicated. Unaffected ubiquitination pattern is indicated with a green box, decreased ubiquitination with a dashed pink box. Results are summarised in H.

The most studied instance for Parkin activation occurs at damaged mitochondria, where activation of Parkin leads to mitophagy (Narendra *et al*, 2008; Matsuda *et al*, 2010). Upon induction of damage, the kinase PTEN-Induced Kinase 1 (PINK1) is stabilised on the outer mitochondrial membrane (OMM) and phosphorylates its primary substrate, ubiquitin, on Ser65 (Kazlauskaite *et al*, 2014b; Koyano *et al*, 2014; Kane *et al*, 2014). Phosphorylated ubiquitin (pUb) is understood to serve for recruitment of Parkin (Okatsu *et al*, 2015), which is normally localised in the cytoplasm, but exhibits nanomolar affinity towards pUb (Kazlauskaite *et al*, 2015; Ordureau *et al*, 2014; Kumar *et al*, 2015). Moreover, PINK1 can also phosphorylate the Ubl of Parkin especially when it is already released from its autoinhibitory site by pUb binding, which leads to further Parkin activation (Kazlauskaite *et al*, 2015; Gladkova *et al*, 2018; Sauvé *et al*, 2018; Kondapalli *et al*, 2012; Shiba-Fukushima *et al*, 2012).

Active Parkin proceeds to ubiquitinate mitochondrial proteins and the resulting ubiquitin chains constitute a signal for mitophagy (Yamano *et al*, 2016). Several cellular studies have suggested the existence of dozens of Parkin substrates and hundreds of target lysines (Ordureau *et al*, 2015; Antico *et al*, 2021; Sarraf *et al*, 2013; Chan *et al*, 2011; Okatsu *et al*, 2012). These include mitochondrial, but also cytoplasmic and even nuclear proteins. The vast number of potential substrates has led to suggestions that Parkin acts in a promiscuous manner when it is brought to proximity of substrates by the presence of pUb (Okatsu *et al*, 2015; Koyano *et al*, 2019; Dunkerley *et al*, 2022; Vranas *et al*, 2022). However, biochemical assays show that Parkin in its active, phosphorylated form (pParkin) ubiquitinates proteins without pUb (Kazlauskaite *et al*, 2014b; Klosowiak *et al*, 2016), which suggests direct interactions between pParkin and its substrates likely exist. An alternative hypothesis is that Parkin interacts with substrates containing ubiquitin-interacting motifs (UIMs) via its Ubl domain (Fallon *et al*, 2006; Chaugule *et al*, 2011; Spratt *et al*, 2013). However, most of the reported Parkin substrates have no UIMs. It has been proposed that the lysines targeted by Parkin are not random and are conserved, although no consensus ubiquitination motif has been found (Klosowiak *et al*, 2016).

Interestingly, Parkin does exhibit preference towards the C-terminal GTPase domain of the mitochondrial GTPase, Miro1 (Klosowiak *et al*, 2016; Dunkerley *et al*, 2022), as compared to its close homolog, Miro2. The difference resides in the presence of a specific lysine K572 in Miro1, which has a particular chemical environment, favourable for ubiquitin conjugation (Klosowiak *et al*, 2016). Importantly, it has been suggested that Parkin acquires specificity when it is phosphorylated, while artificially activated Parkin (for example, containing an N-terminal tag (Burchell *et al*, 2012)) acts indiscriminately (Klosowiak *et al*, 2016). While the molecular basis for preferential ubiquitination of K572 has been described from the Miro1 perspective, there is no explanation for how Parkin might interact with either Miro1 or any of its other substrates, and whether it can indeed directly ubiquitinate all the suggested proteins in its native, non-tagged form.

Here, we evaluate a set of different proteins as Parkin substrates in biochemical assays and show that untagged, full-length Parkin activated with phosphorylation can ubiquitinate many lysine-containing proteins, while it favours Miro1 in a mixture of substrates. We focus on Parkin interaction with its preferred substrate Miro1 and establish an assay to stabilise the Parkin-Miro1 complex. We then characterise the stabilised complex using a series of biochemical and biophysical methods. Most importantly, we identify a substrate-interacting site in Parkin located in the disordered, linker region between Ubl and RING0 domains of Parkin. Upon disruption of Parkin-Miro1 interaction by competition or amino acid substitution, Miro1 ubiquitination by Parkin is compromised. Our findings uncover a substrate recognition mechanism in Parkin and explain previous observations in mitophagy in cells.

## Results

### Active Parkin efficiently ubiquitinates a variety of proteins

Studies report a myriad of potential Parkin substrates, located throughout the cell, including cytoplasmic, mitochondrial, and nuclear proteins (Ordureau *et al*, 2015; Antico *et al*, 2021; Sarraf *et al*, 2013; Chan *et al*, 2011; Okatsu *et al*, 2012). However, these studies often use tagged versions of Parkin which is known to result in artificial Parkin activation (Burchell *et al*, 2012; Matsuda *et al*, 2006; Chaugule *et al*, 2011) and loss of specificity (Klosowiak *et al*, 2016). We therefore wanted to evaluate the promiscuity of untagged, full-length Parkin in biochemical assays. For this, we first looked at ubiquitination of a set of confirmed mitochondrial substrates (Gegg *et al*, 2010; Tanaka *et al*, 2010; Kazlauskaite *et al*, 2014a; Wang *et al*, 2011; Narendra *et al*, 2012; Okatsu *et al*, 2012; Chan *et al*, 2011): Miro1 (HA-tagged, 181-579) (Klosowiak *et al*, 2016), Mitofusin 1 (Mfn1) (6His-tagged, 1-364-GSGSGSGGS-694-741) (Yan *et al*, 2018) and mitoNEET/CISD1 (6His-tagged, 33-108) (Conlan *et al*, 2009) (Suppl. Fig. 1). As previously established, to be active for ubiquitination, Parkin requires the allosteric activator pUb, or phosphorylation at Ser65 of Parkin, or both (Kazlauskaite *et al*, 2014b; Shiba-Fukushima *et al*, 2012; Koyano *et al*, 2014; Kane *et al*, 2014). It has been proposed that Parkin is brought in proximity to its targets via pUb on the mitochondrial membrane (Okatsu *et al*, 2015; Koyano *et al*, 2019; Dunkerley *et al*, 2022; Vranas *et al*, 2022). However, in our assays, we observe that pParkin can ubiquitinate substrates without pUb (Suppl. Fig. 1), consistent with observations by other groups (Kazlauskaite *et al*, 2014a; Klosowiak *et al*, 2016). This suggests that alternate interactions exist between pParkin and its substrates in addition to those driven by pUb recruitment.

Therefore, to simplify our assay set up and focus on direct Parkin-substrate interactions, we performed subsequent assays with pParkin in the absence of pUb. In addition to Miro1, Mfn1 and mitoNEET, we assessed pParkin activity towards pParkin itself, ubiquitin, and typical components of ubiquitination assays such as the E2 enzyme UBE2L3 and the E1 enzyme UBE1 (Fig. 1B). All tested proteins are modified with ubiquitin in the presence of pParkin. In addition, we tested a nuclear protein complex FANCI/FANCD2, not reported as a Parkin substrate in any of the extensive substrate lists, nor as Parkin interactors in the BioGRID database (Oughtred *et al*, 2021). Despite not being a known substrate, the FANCI/FANCD2 complex is also ubiquitinated (Fig. 1B). Finally, we tested a short linear peptide, containing three lysine residues, and find that it is also modified by pParkin *in vitro* (Fig. 1B). Our data confirm that Parkin in its active conformation can modify many proteins that contain solvent-exposed lysines.

### Miro1 is pParkin’s preferred substrate in a mix of proteins

In our assay, pParkin appears to ubiquitinate Miro1 much more extensively than any other tested protein (Fig. 1B, second lane), consistent with reports suggesting Miro1 as the preferred Parkin target (Klosowiak *et al*, 2016; Dunkerley *et al*, 2022; Chan *et al*, 2011). In contrast to the other substrates, there is no detectable unmodified Miro1 and significantly less unconjugated Ub left after 20 minutes of reaction under the conditions used. We therefore hypothesised that pParkin would exhibit preference for Miro1 in a reaction where other substrates are available. To test this hypothesis, we added an excess of Miro1 into each of the reactions with other substrates (Fig. 1C). As expected, the levels of ubiquitination of other substrates decrease as well as levels of available unconjugated Ub (monUb). In contrast, no decrease in Miro1 ubiquitination is observed when we perform a control experiment with excess of Mfn1 or FANCI/D2 (Fig. 1D and E, summarised in Fig. 1F). We then set up a competition assay with various Parkin substrates present in the same reaction: Miro1, Mfn1, mitoNEET, non-conjugatable Ub-6His, inactive Ube2L3 C86A, inactive Parkin C431A and FANCI/FANCD2. Again, pParkin efficiently ubiquitinates each protein (Fig. 1G, lane 3: no excess substrate). Then, we introduced an excess of each substrate at a time, to verify whether pParkin-dependent ubiquitination of Miro1 would be affected. There is no difference in Miro1 ubiquitination in the presence of excess of any of the tested proteins (Fig. 1G, summarised in Fig. 1H). Interestingly, when pParkin is presented with an excess of Miro1, most of Miro1 becomes modified with a few ubiquitins (Fig. 1G, lane 4), rather than extensively polyubiquitinated. This suggests pParkin prefers to ubiquitinate Miro1 itself, rather than to extend Ub chains on Miro1, additionally suggesting that Miro1 is preferred over Ub as a substrate. In addition, when we tested pParkin activity with a limiting concentration of Ub, pParkin only modifies Miro1 and not the other substrates (Fig. 1F, lane 11). Taken together, these data strongly support the hypothesis that Miro1 is the preferred substrate of pParkin, even compared to Ub or Parkin itself, in a mix of substrates.

### Parkin interacts with Miro1 with low affinity

Despite the vast evidence of efficient ubiquitination of various proteins by Parkin, there are no reports of quantifiable, direct interaction between Parkin and any of its targets. In our assays in the absence of pUb, pParkin must interact with its substrates without the action of pUb, likely through transient, low affinity interactions. To capture a Parkin-substrate complex, we chose to investigate the preferred substrate Miro1, reasoning that it is likely to have a stronger interaction with Parkin than any other substrate. Indeed, it has already been suggested that Parkin interacts transiently with Miro1 in cells even in the absence of mitochondrial damage (Safiulina *et al*, 2018). To capture the Parkin-Miro1 complex, we tried several methods, however we were not able to observe a stable complex. In size exclusion chromatography, Parkin and Miro1 elute as separate peaks (Suppl. Fig. 2A). On native PAGE, we do not observe any additional bands as compared to controls (Suppl. Fig. 2B). Isothermal titration calorimetry (ITC) does not detect any Parkin to Miro1 binding (Suppl. Fig. 2C) nor do microscale thermophoresis (MST) experiments with tagged Parkin titrated with Miro1 (Suppl. Fig. 2D) (Dunkerley *et al*, 2022). These data show that the Parkin-Miro1 interaction is weak and cannot be detected by these biophysical methods.

### Parkin-Miro1 complex is detectable upon chemical crosslinking

As we could not detect a stable complex of Parkin and Miro1 through biophysical methods, we wanted to explore whether chemical crosslinking could stabilise the interaction. For this, we used a disuccinimidyl suberate (DSS) crosslinker, which couples primary amines, with a spacer of 11.4 Å. After optimisation, we detect a prominent band corresponding to non-phosphorylated Parkin-Miro1 complex as confirmed by Coomassie staining (Fig. 2A) and Western blot analysis (Fig. 2B). Importantly, a control Parkin-Mfn1 crosslinking reaction produces only a very faint band corresponding to a potential complex, as compared to Parkin-Miro1 crosslinked complex. Our Mfn1 construct has 19 surface-exposed lysines (Fig. 2C) therefore the lack of crosslinking is not due to the unavailability of lysine residues. In contrast, the Parkin-Miro1 crosslinking is easily detectable, indicating a more favourable interaction. The position of the 100 kDa band on SDS-PAGE, compared to the Parkin and Miro1 bands, suggests a 1:1 stoichiometry (Fig. 2A).

**Figure 2:**
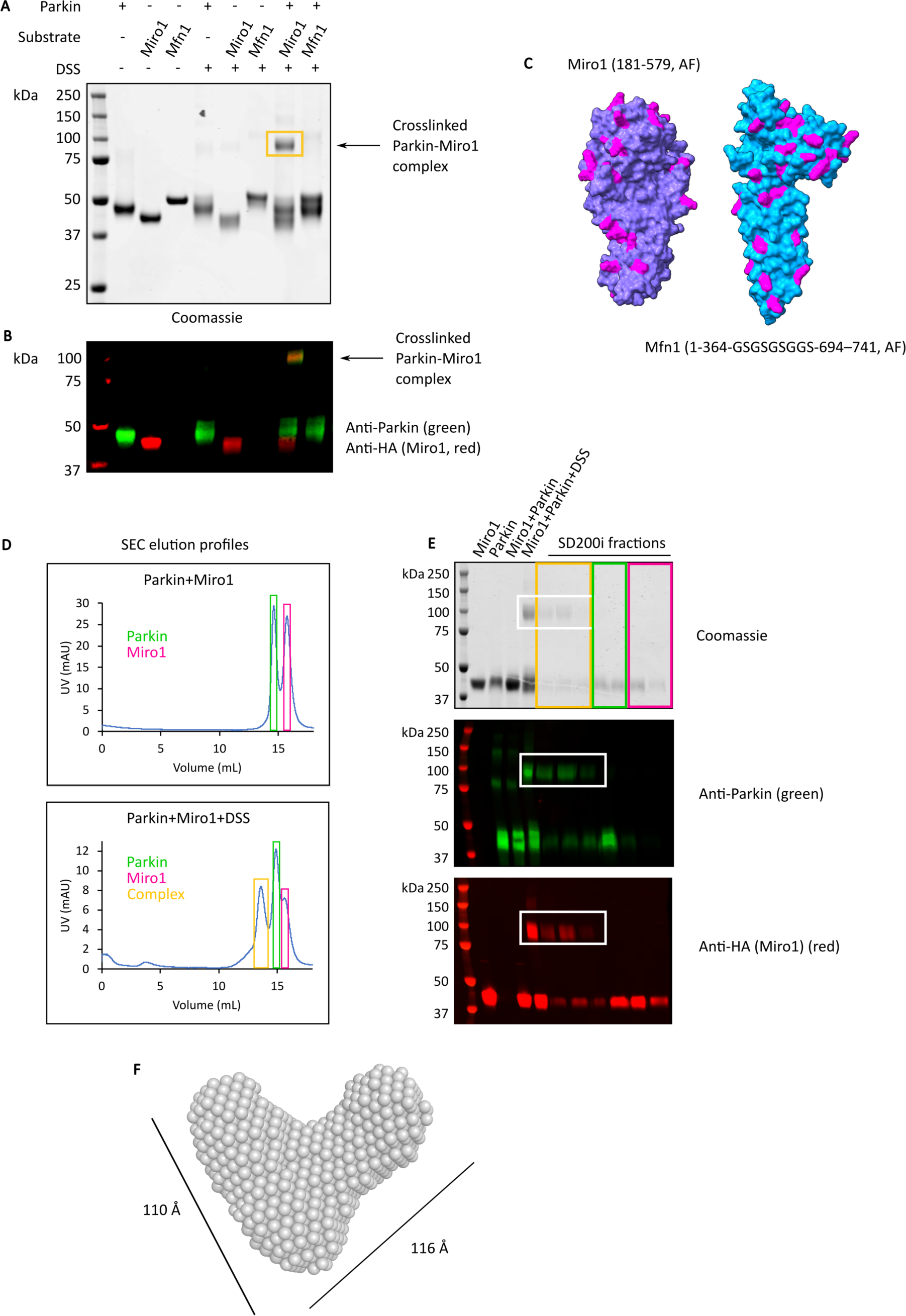
Capture of Parkin-Miro1 complex with a crosslinking assay, isolation and characterisation by SAXS. A: Parkin was incubated with Miro1 or a negative control Mfn1 and the formed complexes were stabilised in a crosslinking reaction by adding DSS where indicated. Controls include each protein separately with and without DSS. After quenching the crosslinking reaction, the samples were resolved by SDS-PAGE and stained with Coomassie or by Western blotting (WB) with anti-Parkin (green) and anti-HA (Miro1) (red) antibodies (B). Band corresponding to the crosslinked 1:1 Parkin-Miro1 complex is indicated with an arrow and yellow box. C: ColabFold-generated surface representation of Miro1 and Mfn1 constructs used in the assay, with exposed lysines coloured in magenta. D: Size exclusion chromatography of crosslinked Parkin-Miro1 complex. Graphs represent elution profiles of the samples without (top) and with the DSS crosslinker (bottom). Third peak corresponding to the complex appears in the SEC run with the crosslinker (marked with yellow box). E: SEC fractions were run on SDS-PAGE and revealed with Coomassie staining or by WB. F: *Ab initio* SAXS model of the crosslinked Parkin-Miro1 complex.

To gain more insight into the size and shape of the Parkin-Miro1 complex in solution, we isolated the crosslinked complex using size exclusion chromatography (SEC) and submitted it to SEC-SAXS analysis (Fig. 2D-F, Suppl. Fig. 3). As controls, we also analysed Parkin and Miro1 separately. The estimated radius of gyration (Rg) for the complex is 39 Å as compared to 28 Å and 28.5 Å determined for autoinhibited Parkin and Miro1 alone, respectively (Suppl. Fig. 3). The Rg values we obtained for Parkin and Miro1 are consistent with the values published previously (Klosowiak *et al*, 2013; Spratt *et al*, 2013). An *ab initio* model of the Parkin-Miro1 complex suggests a V-shaped conformation, with the length of the branches estimated at around 110 Å and 116 Å (Fig. 2F), likely corresponding to the two crosslinked proteins linked near their termini, with the other extremities pointing outwards. These data indicate an interaction between Parkin and Miro1 that is stabilised by chemical crosslinking.

### Identification of Parkin-Miro1 interaction site

After establishing the crosslinking assay to capture a Parkin-Miro1 complex, we sought to identify the regions of Parkin that are important for the interaction with Miro1. For this, we performed crosslinking reactions with various Parkin deletion constructs, devoid of Ubl, linker, Ubl and linker, or Rcat (Fig. 3A). Interestingly, only the constructs without linker exhibit decreased crosslinking efficiency with Miro1, suggesting that the linker element of Parkin is involved in the interaction. Of note, we have previously observed that Parkin’s ability to ubiquitinate Miro1 is compromised when the linker region is deleted (Kumar *et al*, 2015). No decrease in complex formation is observed with the deltaRcat variant, which suggests the interaction of Rcat with the substrate, although likely necessary for the ubiquitination activity (Klosowiak *et al*, 2016), does not contribute to the overall stability of the complex. Interestingly, when we perform the crosslinking reaction with phosphorylated Parkin, we do not observe increased intensity of the band corresponding to the crosslinked complex. This indicates that crosslinking of Parkin and Miro1 is not dependent on the activation state of Parkin, consistent with previous cellular observations (Safiulina *et al*, 2018).

**Figure 3:**
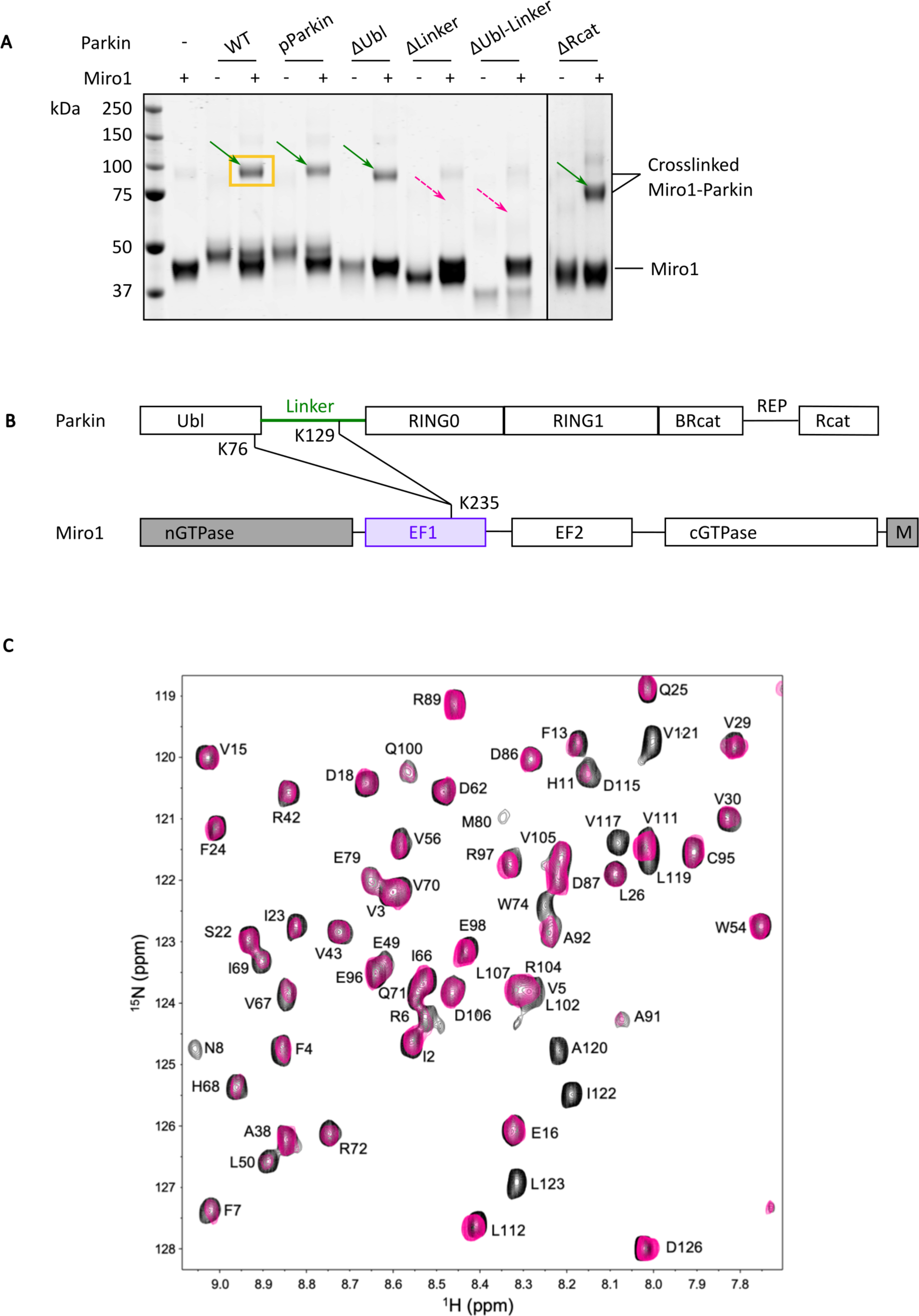
Identification of Miro1-interacting region in Parkin using crosslinking assay and NMR spectroscopy. A: Series of Parkin mutants with various domains deleted were assayed in the crosslinking assay for their ability to form a crosslinkable complex with Miro1. Green arrows indicate bands corresponding to crosslinked complex while dashed pink arrows point to areas where bands for crosslinked complex were not detected. B: Schematic of Parkin and Miro1 domains with indicated crosslinked lysines as identified by mass spectrometry. Parkin construct is full length, while Miro1 construct (residues 181-579) does not have the N-terminal GTPase domain nor the C-terminal transmembrane domain (grey). C: Interaction of Ubl1-126 and Miro1180-582 using NMR spectroscopy. Overlay of regions of the 1H-15N HSQC spectra of 15N Ubl1-126 alone (black contours) and 15N-labelled Ubl1-126 with one equivalent of unlabelled Miro1180-582 (pink contours). Residues are labelled according to their one-letter amino acid code and number. G114 and S116 are not shown in this region.

To investigate the details of the interaction, we analysed the crosslinked full-length Parkin-Miro1 Coomassie-stained band by mass spectrometry. We find that lysines 76 and 129 in Parkin, located in the C-terminus of the Ubl domain and in the linker region respectively, are uniquely crosslinked to lysine 235 in the EF1 domain in Miro1 (Fig. 3B, Suppl. Fig. 4). This further confirms that the linker region in Parkin is involved in the interaction with Miro1, while the Ubl domain remains in crosslinkable proximity (<30 Å, (Merkley *et al*, 2014)) to the interaction site.

To further explore the Miro1 interaction region in Parkin, we used NMR spectroscopy. We titrated the EF1-EF2-cGTPase region of Miro1 (Klosowiak *et al*, 2016) (residues 180-582, Miro^180-582^) into an 15N-labelled Ubl-linker construct (Ubl^1-126^) from Parkin. In the absence of Miro^180-582^ the ^1^H-^15^N HSQC spectrum of the Ubl^1-126^ displays well-resolved signals from both the folded Ubl domain and the linker region (Fig. 3C). Notably, most signals for the linker region between M80-D126 fall within 8.0-8.5 ppm indicative of a disordered protein structure. Upon addition of Miro^180-582^ signals from residues D115-V117 and L119-L123 broaden or shift beyond recognition. This observation is typical of a weaker interaction between the two proteins on the intermediate time scale, in agreement with our biophysical experiments. In contrast, all signals from the folded Ubl domain (I2-V70) retain similar intensities and positions in the spectrum. These observations are consistent with binding of the D115-L123 region in Parkin to the 44 kDa fragment of Miro1, with little or no involvement of the Ubl domain, in agreement with our observations from the crosslinking assay.

### AlphaFold prediction of Parkin-Miro1 complex indicates a confident region of interaction

To further characterise the elusive Parkin-Miro1 interaction, we employed AlphaFold using ColabFold (Jumper *et al*, 2021; Mirdita *et al*, 2022). AlphaFold predicts a model for Parkin-Miro1 complex with low confidence with regard to the relative orientation of Parkin and Miro1 domains, as indicated by the Predicted Aligned Error (PAE) plot (Fig. 4A, B,). Importantly however, there is one region of Parkin predicted as interacting with high confidence with Miro1. It is positioned within the linker of Parkin, as identified in our crosslinking-mass spectrometry experiments, and, more specifically, it overlaps with the region identified from NMR experiments. It corresponds to residues 115-124 (DSVGLAVILHT) in Parkin, which we named the Substrate Targeting Region (STR) (Fig. 4A-C). The STR is conserved from fish to humans (Fig. 4C) and includes the hydrophobic ‘GLAVIL motif’ (Fig. 4C) positioned downstream to the conserved ‘ACT element’ (Gladkova *et al*, 2018). The STR is predicted by AlphaFold to thread through the EF1 domain of Miro1, with the hydrophobic residues in the STR (V117, L119, A120, V121, I122, L123) binding to a deep hydrophobic pocket in Miro1 (Fig. 4F, G). The hydrophobic interactions are gated by a triad of charged residues D115, D126 and R128 with the potential to form salt bridges with Miro1 residues R263, R265 and D228, respectively (Fig. 4F). AlphaFold contacts, corresponding to pairs of residues that coevolved in the two interacting proteins, extend to 15, mostly hydrophobic, residues in the Miro1 pocket, paired with Parkin residues in the 115-124 stretch (Fig. 4D, E). Alternative AlphaFold models indicate the position of the STR in Parkin relative to Miro1 remains consistent with high confidence scores for the interaction (Suppl. Fig. 5A-C).

**Figure 4:**
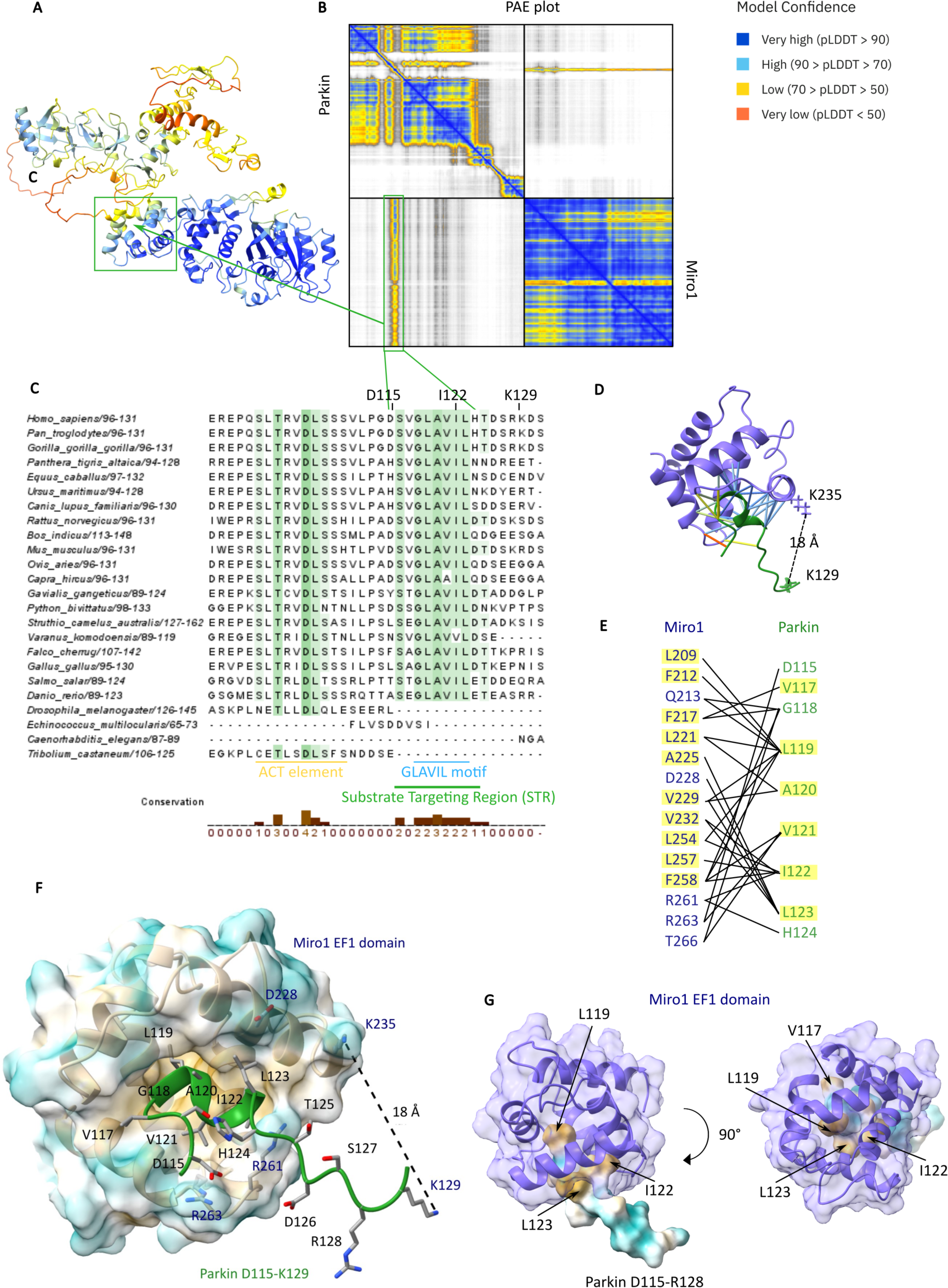
High-confidence AlphaFold prediction of Parkin-Miro1 interface which is driven by hydrophobic interactions. A: Ribbon representation of top-ranked AlphaFold model of Parkin-Miro1 complex, coloured by model confidence (pLDDT) as indicated in the legend. B: Predicted Aligned Error (PAE) plot reveals a stretch of residues in Parkin linker predicted with high confidence to bind to Miro1. Green boxes highlight the region of interest. C: Sequence alignment of Parkin linker region (residues E98-S131 in *Homo sapiens*) reveals two patches of residues with higher conservation: the ACT motif and the Miro1 interacting, Substrate Targeting Region (STR), encompassing the GLAVIL motif. D: Close-up on the AlphaFold coevolution contacts between the EF1 domain in Miro1 (purple) and Parkin linker region (residues D115-K129, green). The lysines K129 (Parkin) and K221 (Miro1) which were identified as crosslinked by MS and the predicted distance between them are indicated. E: List of the coevolved residue pairs in Miro1 and Parkin as illustrated in D. Hydrophobic residues are highlighted In yellow. F: Hydrophobic surface representation (yellow: hydrophobic residues, blue: hydrophilic residues) of Parkin binding pocket in the Miro1 EF1 domain with green ribbon representing the linker region in Parkin (D115-R128) which contains the Miro1 interacting region. Residues of interest are represented as sticks. G: View of the Miro1 EF1 domain (purple ribbon and surface representation) and Parkin D115-R128 region represented as hydrophobic surface, with key hydrophobic residues in Parkin indicated by arrows.

Since we used a Miro1 construct which has its N-terminal GTPase domain deleted, we wondered whether full-length Miro1 would be predicted to bind to Parkin differently. However, the AlphaFold model for the full-length Miro1 in complex with Parkin closely resembles the one with our Miro1 construct (Suppl. Fig. 5D-E), with the N-terminal domain flexibly connected to the remaining Miro1 core. This prediction suggests that the N-terminal GTPase does not contribute to the interaction with Parkin.

We noticed that in the AlphaFold model, the position of the crosslinked K76 residue in Parkin, (Fig. 3B) is predicted to be over 30 Å away from K235 in Miro1, suggesting that the Ubl might be mobile with respect to Miro1. Ubl mobility has been previously observed, especially for pUbl (Aguirre *et al*, 2017; Condos *et al*, 2018; Gladkova *et al*, 2018). Indeed, the prediction for the position of Ubl and other Parkin domains relative to Miro1 is of low confidence in our AlphaFold models and cannot be confirmed by our SAXS analysis either due to the limited resolution of SAXS, although the *ab initio* SAXS model overlaps well with the AlphaFold model (Suppl. Fig. 5F). Importantly however, the Parkin K129 – Miro1 K235 residue pair detected in our crosslinking-mass spectrometry experiment is predicted by AlphaFold to be 18 Å away (Fig. 4D), which is well within the crosslinkable distance, supporting the proposed position of the interacting region.

### Substitutions in the interacting region disrupt Miro1 ubiquitination by Parkin

With STR identified as the Miro1-interacting region in Parkin, we hypothesised that substitutions in that region would alter Parkin-Miro1 interaction and Parkin-dependent ubiquitination of Miro1. To test this hypothesis, we chose the residue I122 in Parkin, as the residue making several coevolution ‘contacts’ with Miro1’s hydrophobic pocket, based on AlphaFold predictions (Fig. 4E) and one of the hydrophobic residues within the Miro1 EF1 pocket, L221, coevolved with a different set of Parkin residues (Fig. 5A). We first assayed Parkin I122Y and Miro1 L221R in our crosslinking assay. Strikingly, these single amino acid substitutions substantially decrease formation of the crosslinked complex (Fig. 5B). Next, we tested the binding of a fluorescently labelled STR-containing peptide (Cy5-STR) to Miro1 by MST. While we are unable to reach binding saturation and conclude binding affinity due to the limitations of the method for studying low affinity interactions, we can nonetheless observe differences in the binding profile of WT versus L221R Miro1, with the substituted Miro1 binding compromised (Fig. 5C).

**Figure 5:**
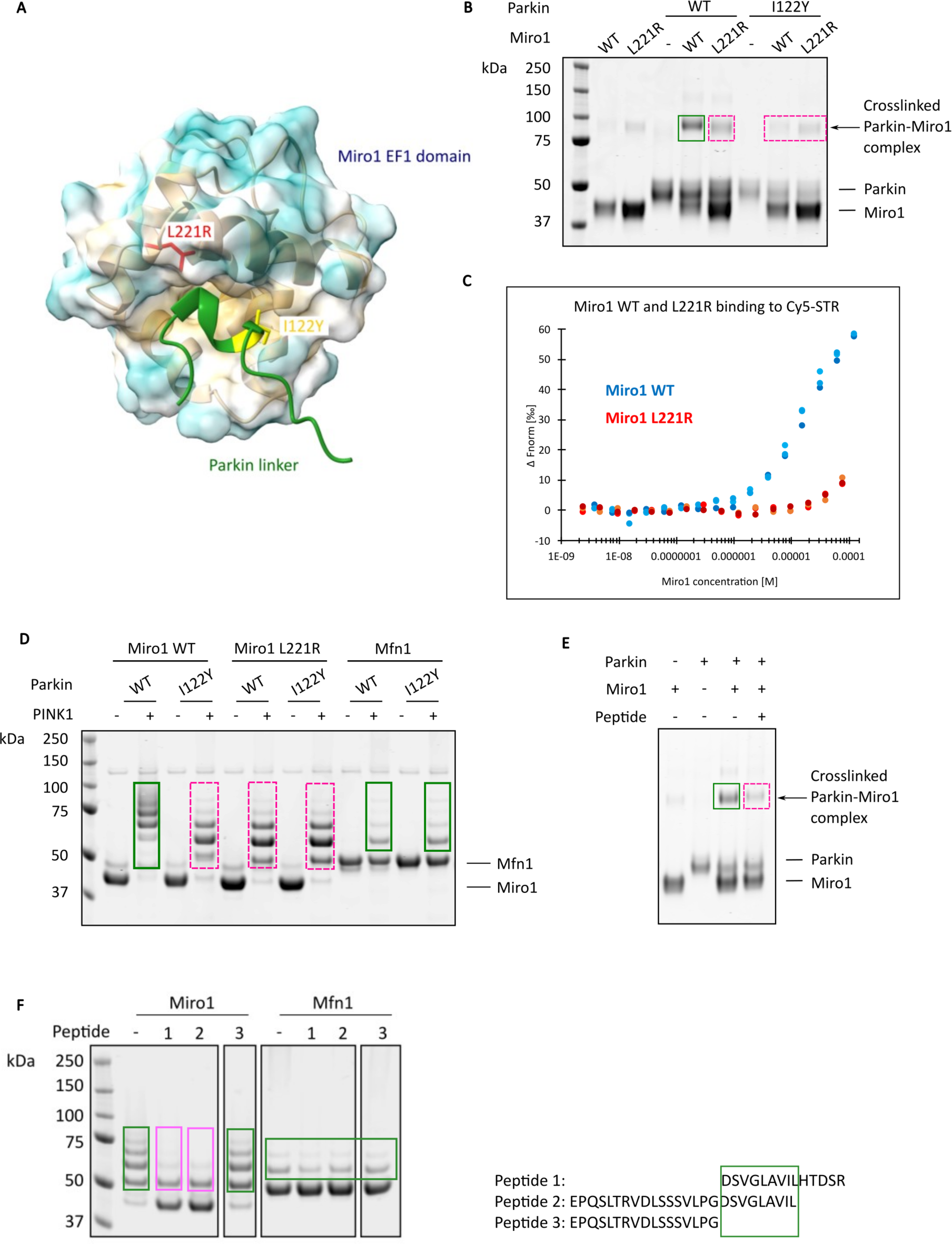
Substitutions in the STR and Miro1, and synthetic STR-containing peptides disrupt Miro1 binding and ubiquitination by Parkin. A: Position of the mutated residues (red and yellow) in the EF1 domain of Miro1 (surface) and in Parkin linker (green ribbon) are indicated. B: Crosslinking assay with mutated Miro1 and/or Parkin shows decrease in Miro1-Parkin complex formation (dashed pink boxes) as compared to wild-type Parkin and Miro1 (green box). C: Binding of fluorescent synthetic peptide from Parkin linker region containing STR (Cy5-STR) to Miro1 WT (blue) and L221R mutant (red) was assayed using MST. Affinity of Miro1 L221R mutant to Cy5-STR was decreased, although affinities could not be calculated. D: Miro1 WT and L221R mutant or Mfn1 (control) as indicated were subject to ubiquitination assay with Parkin WT and/or I122Y mutant. Decreased Miro1 ubiquitination as compared to WT (green) was marked with dashed pink boxes. E: Crosslinking assay with Parkin and Miro1 in the presence of the STR-containing peptide. F: Miro1 ubiquitination assay with a series of synthetic STR-containing peptides. Dashed pink boxes indicate decreased Miro1 ubiquitination. The sequences of corresponding peptides are indicated on the right. Ubiquitination assay with Mfn1 was used for comparison of Parkin activity in the presence of the peptides. Green box indicates the residue stretch in the STR-containing peptide, responsible for decreased Miro1 ubiquitination by Parkin.

Next we wanted to verify whether Parkin ubiquitination of Miro1 is indeed dependent on the STR interaction with the hydrophobic pocket in the Miro1 EF1 domain. We therefore set up a ubiquitination assay with mutated Parkin and Miro1. Substitution of I122Y in Parkin indeed causes a significant decrease in Miro1 ubiquitination (Fig. 5D), suggesting that a single amino acid change in the STR is sufficient to disrupt Parkin ability to ubiquitinate Miro1. Importantly, the Parkin I122Y protein does not exhibit any change in the levels of ubiquitination of a control substrate Mfn1, indicating that the I122Y mutation affects Miro1 ubiquitination specifically, but does not influence the Parkin activity towards substrates in general. As for Miro1, the L221R variant is not ubiquitinated as efficiently as the WT. Importantly, when both Parkin I122Y and Miro1 L221R mutants were tested in the same reaction, the decrease of Parkin activity towards Miro1 is not significantly more pronounced as compared to reaction with either Parkin or Miro1 mutant. The fact that the effects of the mutations are not additive indicates the mutagenesis affects the same interaction site, providing further support for the interaction between STR and the hydrophobic EF1 pocket in Miro1.

### Synthetic STR-containing peptides compete for Miro1 binding

We reasoned that if the STR is the main site of Parkin interaction with Miro1, addition of a synthetic peptide that contains the STR sequence should compete for the Parkin binding site on Miro1. To test this, we synthesised an STR-containing peptide (Parkin residues D115-R128) and added it to our crosslinking and activity assays. As expected, we observe a decrease in Parkin-Miro1 binding upon adding excess STR peptide as detected by crosslinking (Fig. 5E). We also observe a decrease in Miro1 ubiquitination by Parkin in an activity assay (Fig. 5F, peptide 1), further supporting the hypothesis that the excess of synthetic STR peptide interferes with Parkin-Miro1 binding and subsequent Miro1 ubiquitination. At high concentrations of peptide, we also observe a slight decrease in Parkin activity towards Mfn1, which suggests STR may have some additional role in Parkin activity regulation or participate in targeting of Mfn1, although to a much lower extent than in targeting Miro1 (Suppl. Fig. 6A). To verify that the inhibition is not due to chemical contamination from the peptide synthesis, we tested another unrelated peptide acquired from the same source. We do not observe any effect on Miro1 ubiquitination with this control peptide (Suppl. Fig. 6B).

To further dissect the region in the STR peptide sequence which is important for Miro1 binding and its subsequent ubiquitination, we tested a series of STR-containing peptides covering Parkin residues E98-R128 (also containing the ACT element and the GLAVIL motif (Gladkova *et al*, 2018)) in Miro1 activity assays (Fig. 5F). Only peptides which contain residues D115-L123 (DSVGLAVIL) are inhibitory, and the presence of the ACT element residues and before (E98-G114) does not have any detectable effect.

## Discussion

Several studies have reported a surprisingly large number of proteins identified as ubiquitinated by Parkin under mitophagy-inducing conditions (Antico *et al*, 2021; Ordureau *et al*, 2014; Sarraf *et al*, 2013). However, it has remained unknown whether all identified proteins are direct Parkin substrates, and if yes, how Parkin recognises and ubiquitinates such a diverse range of proteins. Moreover, no substrate recognition motif has been found in Parkin. Here, we show that full-length, untagged Parkin activated by PINK1 acts in a promiscuous manner and can directly ubiquitinate many different proteins in a biochemical assay. We establish that Parkin favours Miro1 as a substrate in a mix of proteins, including ubiquitin and Parkin itself, suggesting that Parkin does have some level of specificity. Further, despite the challenges encountered due to the low affinity of the interaction, through a series of biochemical approaches, we demonstrate the existence of an elusive Parkin-substrate interaction. We show that Parkin binds to Miro1 via a short hydrophobic region in the mostly disordered linker between Ubl and the RING0 domain of Parkin. Our findings highlight the functional importance of this moderately conserved region of Parkin, so far ‘invisible’ in structures determined using X-ray crystallography due to its disordered nature, with the potential for conditional folding that we predict computationally. Knowing that Parkin can interact with Miro1, it is now tempting to speculate that Parkin could interact with other, not yet investigated proteins via the same or a different interaction site. On the other side, the hydrophobic Miro1 pocket has been proposed to accommodate various other partners besides Parkin (Covill-Cooke *et al*, 2024), which suggests Parkin may be competing with other proteins such as cytoskeletal adaptors CENPF, Trak and MYO19 for binding to Miro1 in cells.

The Parkin preference for targeting Miro1 as compared to its close homolog Miro2 was previously described in detail and attributed to the specific chemical environment of the favoured lysine in the cGTPase domain of Miro1 (Klosowiak *et al*, 2016). While the cGTPase domain on its own is sufficient and necessary for Miro1 ubiquitination by Parkin, the presence of the EF1 domain leads to a slight increase in Miro1 and Miro2 ubiquitination, which we can now explain by increase in affinity resulting from Parkin binding to EF1. What emerges is a two-step Miro1 preference by Parkin: initially driven by the EF1-Parkin binding, and upon Parkin activation, by targeting a specific lysine on the cGTPase domain of Miro1.

Importantly, our results suggest that Parkin can interact with Miro1 independently of its activation state, as Parkin phosphorylation does not detectably increase its interaction with Miro1 based on crosslinking data. Together with the two-step preference for Miro1, this would suggest that Parkin binding to a substrate and its ubiquitination activity are uncoupled, and, at least in this specific case, Parkin activity is not induced by substrate binding. Interestingly, Parkin interaction with Miro1 has been previously observed in cellular studies, where Parkin translocation to mitochondria upon mitochondrial damage was dependent on Miro1/2 expression but occurred independently of PINK1 expression and consequently, of Parkin activation state (Safiulina *et al*, 2018). This suggests that Miro1 serves as a mitochondrial acceptor for Parkin even before activation of the PINK1/Parkin cascade, which generates pUb chains considered to be the main Parkin receptors (Okatsu *et al*, 2015; Vranas *et al*, 2022; Dunkerley *et al*, 2022; Koyano *et al*), and does not get ubiquitinated until mitochondria are damaged and Parkin is activated. Consequently, a small pool of Parkin is constantly present at the mitochondrial membrane, which has been indeed observed in cells (Narendra *et al*, 2008; Darios *et al*, 2003; Safiulina *et al*, 2018; Shlevkov *et al*). While the significance of this is unclear, we speculate that low levels of Parkin at the mitochondrial membrane, readily available upon PINK1-induced activation for a rapid ubiquitination of substrates and further Parkin recruitment through the feed-forward mechanism (Ordureau *et al*, 2014), allow for a rapid response to mitochondrial damage. This may be important in dopaminergic neurons that are susceptible to mitochondrial stressors and where fast mitochondrial turnover is critical to prevent neurodegeneration (Haddad & Nakamura, 2015). Of note, in a very recent cellular study focusing on Miro1 interactors, including Parkin, which may bind to the hydrophobic pocket in the EF1 domain, Parkin translocation was diminished upon L119A mutation, supporting the importance of the Parkin Miro1-interacting site in mitophagy. While no pathogenic mutations in that site have been reported so far, ClinVar database lists two missense mutations of uncertain significance (D115G and G118A), identified in patients with autosomal recessive juvenile Parkinson disease, cancer and inborn genetic diseases (Rehm *et al*, 2015). Our work brings a new perspective on Parkin specificity towards substrates and its involvement in mitophagy and it may influence directions for therapeutic avenues in neurological disorders where mitophagy is affected, such as in Parkinson’s disease.

## Methods

### Protein expression and purification

His6-Mfn1 (1-364-GSGSGSGGS-694-741), HA-Miro1 (181-579), His6-mitoNEET (33-108), FancI/D2, GST-PINK1 (126-C), UBE2L3, pUb and Ub-MES were purified as previously described (Chaugule *et al*; Kumar *et al*, 2015; Conlan *et al*, 2009; Yan *et al*, 2018; El Oualid *et al*, 2010).

Parkin was produced using modified protocols (Chaugule *et al*, 2011). Briefly, codon-optimised His6-Smt3-Parkin constructs were expressed in *E. coli* BL21(DE3) Rosetta cells at 37°C to an OD600 of 0.7 in 2xYT media supplemented with 0.5 mM ZnCl2. Expression was induced at 16°C with 300 μM IPTG for 18 h. After Ni^2+^NTA affinity purification, His6-Smt3 tag was cleaved with Ulp1 protease overnight at 4°C, followed by ResourceQ ion exchange chromatography and Superdex 75 10/300 size exclusion chromatography. For pParkin, after the first purification step with Ni^2+^NTA affinity, Parkin was eluted and phosphorylated with PhPINK^126-C^ while dialysing in phosphorylation buffer (5 mM ATP, 5 mM MgCl2, 50 mM NaCl, 50 mM Tris-HCl pH 8, 1 mM DTT) for 1 h at 20°C. pParkin was then re-bound to Ni^2+^NTA resin and further purified as for non-phosphorylated Parkin.

HA-Miro1^181-579^ was purified as follows: *E. coli* BL21 (DE3) cells were grown in 2xYT media until OD600 0.7, cooled down to 18°C and induced with 300 μM IPTG for 18 h. Cells were lysed by sonication in lysis buffer (PBS with additional 200 mM NaCl, 0.4 mM TCEP, 5% glycerol) supplemented with 4 mM MgCl2, protease inhibitor tablets (Pierce, Fisher Scientific) and DNAse I (Merck). After lysate clearing by centrifugation, GST-tagged Miro1 construct was purified on GSH agarose and the tag was cleaved using 3C protease overnight at 4°C. Miro1 was then concentrated and purified on Superdex 75 10/300 size exclusion chromatography column.

Parkin and Miro1 point mutants were generated by site-directed mutagenesis, confirmed by sequencing and purified as above.

### Commercial reagents

STR-containing peptides were synthesised by Genosphere Biotechnologies with >95% purity as assessed by HPLC and mass spectrometry, and dissolved in 100% DMSO. Ubiquitin was purchased from Biotechne (U-100H) as lyophilised powder and dissolved in 100 mM NaCl, 20 mM HEPES pH 7.4. DSS crosslinker was acquired from Fisher Scientific and dissolved in 100% DMSO.

### Ubiquitination assays

For substrate ubiquitination assays, each substrate at 2 µM (besides FancI/D2 complex, which was at 0.4 µM due to a substantially larger number of lysines available for ubiquitination as compared to the other substrates) was mixed with 40 µM Ub, 0.5 µM E1, 2 µM Ube2L3, 0.5 µM pParkin, and an excess of another substrate was added as indicated at 10 µM, in ubiquitination buffer (50 mM Tris-HCl pH 7.5, 100 mM NaCl, 5% glycerol, 5 mM MgCl2). For the low concentration Ub reaction, 2 µM of ubiquitin was used instead of 40 µM. Reactions were supplemented with 5 mM ATP where indicated and incubated for 20 min at 37°C.

For Miro1 ubiquitination reactions with mutated Parkin and Miro1, 4 µM of Miro1 (or Mfn1 as control) was incubated with 20 µM Ub, 0.1 µM E1, 0.5 µM Ube2L3, 0.5 µM Parkin or pParkin as indicated and 50 nM PINK1 where indicated, in ubiquitination buffer and ATP as above. Reactions were stopped after 45 min at 37°C.

For Miro1/Mfn1 reactions with synthetic peptides, 4 µM of Miro1 or Mfn1 was incubated with 20 µM of the reactive ubiquitin Ub-MES (to eliminate potential interactions of the peptide with E1 or E2) and 0.5 µM pParkin in ubiquitination buffer and ATP as above. 100 µM of peptide was added unless specified otherwise, to a final of 2% DMSO, and the reaction was incubated for 1 h at 37°C.

The reactions were stopped by adding NuPAGE^TM^ 4x LDS loading buffer (Novex) and 167 µM DTT prior to being resolved on NuPAGE^TM^ LDS 4-12% Bis-Tris gels (Novex). Gels were stained with a Coomassie-based SimplyBlue^TM^ SafeStain (Invitrogen) and scanned using LI-COR® Odyssey Clx Infrared Imaging System at 700 nm wavelength.

### Crosslinking and mass spectrometry

3 µM of Parkin constructs were mixed with 3 µM HA-Miro1^181-579^ in 50 mM HEPES pH 7.4, 150 mM NaCl, 0.4 mM TCEP, 5% glycerol in a 10 µL volume and preincubated for 5 min at room temperature (20°C). Then 0.3 mM of disuccinimidyl suberate (DSS) crosslinker (Thermo Scientific™) in DMSO was added and the reaction was incubated for further 45 min. Reaction was stopped by adding 50 mM Tris-HCl pH 8. The samples were resolved on a NuPAGE^TM^ LDS 4-12% Bis-Tris gels (Novex) under reducing conditions and Coomassie-stained. Bands corresponding to expected Parkin-Miro1 complex size (∼100 kDa) were excised and sent for mass spectrometry (BSRC Mass Spectrometry & Proteomics Facility, University of St Andrews, St Andrews, UK). The samples were reduced, alkylated and trypsin-digested at the Facility, and the resulting peptides were analysed by LC-MSMS on an Orbitrap Fusion™ Lumos™ mass spectrometer (ThermoFisher Scientific) using CID+ETD activation. The crosslinked peptides were identified using MeroX 2.0.1.4 software (Götze *et al*, 2015, 2012) with default settings (1% false discovery rate, 10% prescore intensity, 3 missed cleavages allowed) with following changes: phosphorylation included as post translational modification, BS3/DSS crosslinker, specificity site: lysines only.

### SEC and SEC-SAXS

For the large scale crosslinking and purification of the Parkin-Miro1 complex, the crosslinking reaction was performed as described above but in 10 mL final volume, in 50 mM HEPES pH 7.4, 150 mM NaCl, with 0.2 mM DSS. After incubation and quenching, the sample was concentrated and purified on a Superdex 200 Increase 10/300 size exclusion column at 0.25 mL/min. Fractions from the peak corresponding to the Parkin-Miro1 complex were collected, analysed by SDS-PAGE and Western blotting, concentrated and flash-frozen in liquid nitrogen.

The samples were shipped to Diamond Light Source, beamline B21. Samples were applied onto a Superdex 200 Increase 3.2 column equilibrated in a buffer comprising 150 mM NaCl, 50 mM HEPES pH 7.4, 0.4 mM TCEP, and 5% glycerol, at 0.16 mL/min, before exposure to x-rays as part of the standard setup at the beamline. Data were analysed using ScÅtter version IV.j (available from beamline B21 at Diamond Light Source). *Ab initio* models were calculated by 21 independent runs of DAMMIF (Franke & Svergun, 2009) followed by averaging and filtering using DAMAVER (Volkov & Svergun, 2003).

### AlphaFold predictions using ColabFold and model visualisations

ColabFold (Mirdita *et al*, 2022; Jumper *et al*, 2021) version 1.5.2.20231005 has been used through the MVLS Advanced Research System (MARS) computing platform at the University of Glasgow with the following parameters: number of recycles: 12, number of models: 5, with amber relaxation. Models were visualised using PyMOL (The PyMOL Molecular Graphics System, Version 3.0 Schrödinger, LLC) or with UCSF ChimeraX, developed by the Resource for Biocomputing, Visualization, and Informatics at the University of California, San Francisco, with support from National Institutes of Health R01-GM129325 and the Office of Cyber Infrastructure and Computational Biology, National Institute of Allergy and Infectious Diseases (Goddard *et al*, 2018).

### NMR Experiments

The autoinhibitory Ubl domain and linker region of human parkin (residues 1-126, Ubl^1-126^) was created by inserting a stop codon in full-length parkin. Ubl^1-126^ was expressed as a His6-SUMO fusion protein in *E. coli* BL21(DE3) cells using M9 minimal media supplemented with ^15^NH4Cl (1.0 g/L) for NMR experiments purified as previously reported ((Aguirre *et al*, 2017; Chaugule *et al*, 2011)). Samples of 15N-labelled Ubl^1-126^ (140 µM) were prepared in 50 mM Tris, 150 mM NaCl, 250 µM TCEP (pH 7.5) with 10% (v/v) D2O. Imidazole was included as a pH indicator and 4,4-dimethyl-4-silapentane-1-sulfonic acid (DSS) as a direct internal ^1^H reference. ^1^H-^15^N HSQC spectra were acquired at 25°C on a Bruker Avance Neo 600 MHz NMR spectrometer equipped with a triple resonance probe using z-field gradients (BioCORE Facility, Western University). Spectra were collected in the absence and presence of an equimolar amount of Miro1^180-582^. Data were processed using NMRPipe (Delaglio *et al*, 1995), visualized using NMRViewJ (Johnson & Blevins, 1994) and assigned using previously reported chemical shift data (Aguirre *et al*, 2017).

### MST

For Parkin-Miro1 binding experiments, His6-Smt3-tagged Parkin in 50 mM HEPES, 150 mM NaCl, 5% glycerol with 0.1% Tween20 was labelled with Monolith His-Tag Labelling Kit RED-tris-NTA 2nd Generation (NanoTemper) according to the manufacturer’s protocol. Miro1 was prepared in a series of concentrations in 50 mM HEPES, 300 mM NaCl, 5% glycerol and titrated into Parkin samples in 1:1 volume ratio. Measurements were performed at 18-22°C on a Monolith NT.115 instrument (NanoTemper Technologies) using the red channel, using 40% excitation power and 40% MST power. For the STR-containing peptide binding experiments to Miro1, fluorescent Cy5-STR peptide at the final concentration of 20 nM in PBS with 2.5% DMSO, 0.05% Tween20 and 0.5 mg/mL bovine serum albumin was titrated with Miro1 WT or Miro1 L221R. Measurements were performed with 20% excitation power and 40% MST power. Data was analysed using the MO.Affinity Analysis v2.3 (NanoTemper Technologies).

## Acknowledgements

The authors would like to thank the BSRC Mass Spectrometry & Proteomics Facility, University of St Andrews, St Andrews, UK, for the mass spectrometry experiments, June Southall from the Integrated Protein Analysis Facility, Shared Research Facilities, University of Glasgow, for help with MST and ITC experiments, and Diamond Light Source for beamtime for the SEC-SAXS experiments (BAG allocation mx28516). This research was supported through grants from the Wellcome Trust (209347/Z/17/Z, HW) and Canadian Institutes of Health Research (PJT 166019, GSS).

