## Supplementary figures for "A substrate-interacting region of Parkin directs ubiquitination of the mitochondrial GTPase Miro1"

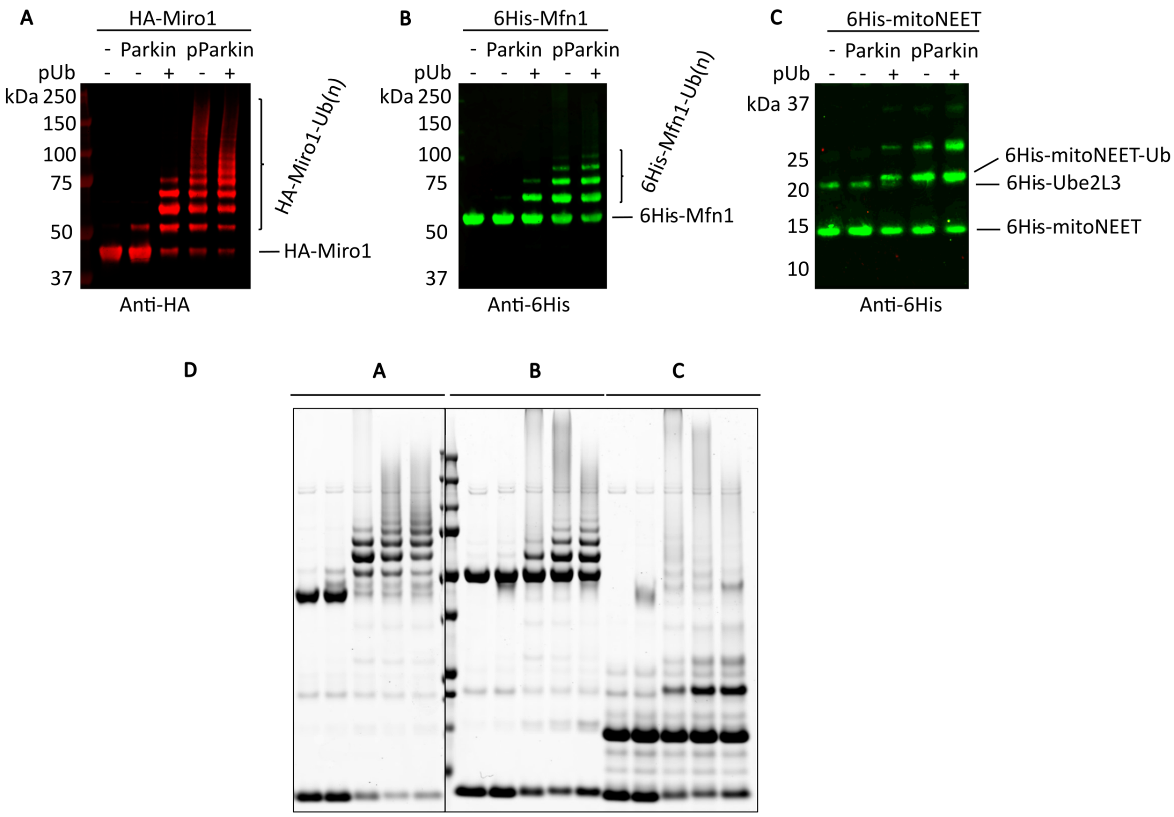


**Supplementary Figure 1: Ubiquitination assays of Parkin mitochondrial substrates.** Substrates Miro1 (A), Mfn1 (B) and MitoNEET (C) were submitted to Parkin ubiquitination assays in the presence or absence of pUb and with Parkin or pParkin, as indicated. Reactions were resolved by SDS-PAGE and revealed with WB. D: Corresponding Coomassie-stained gels.

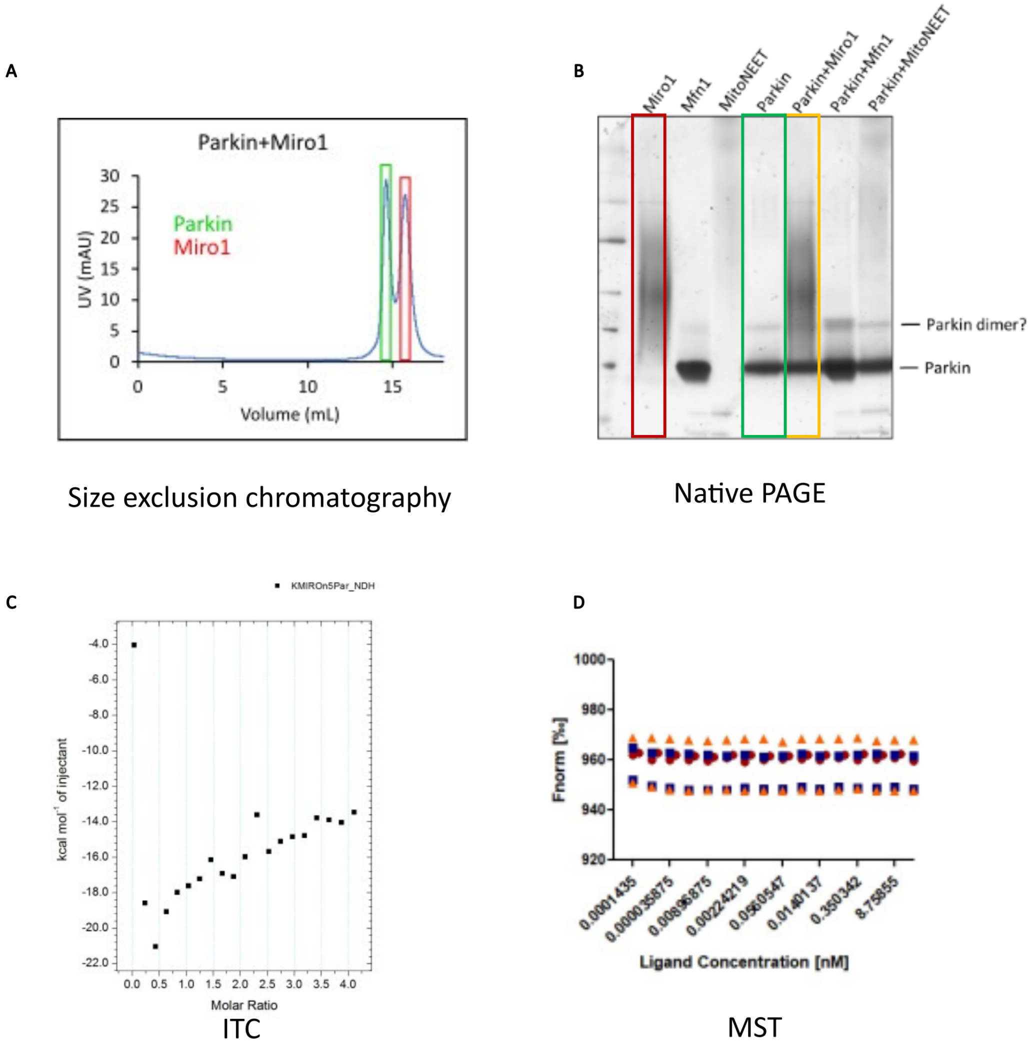


**Supplementary Figure 2: Results from attempts using various methods to capture Parkin-Miro1 complex**. A: SEC, B: native PAGE, C: ITC, D: MST.

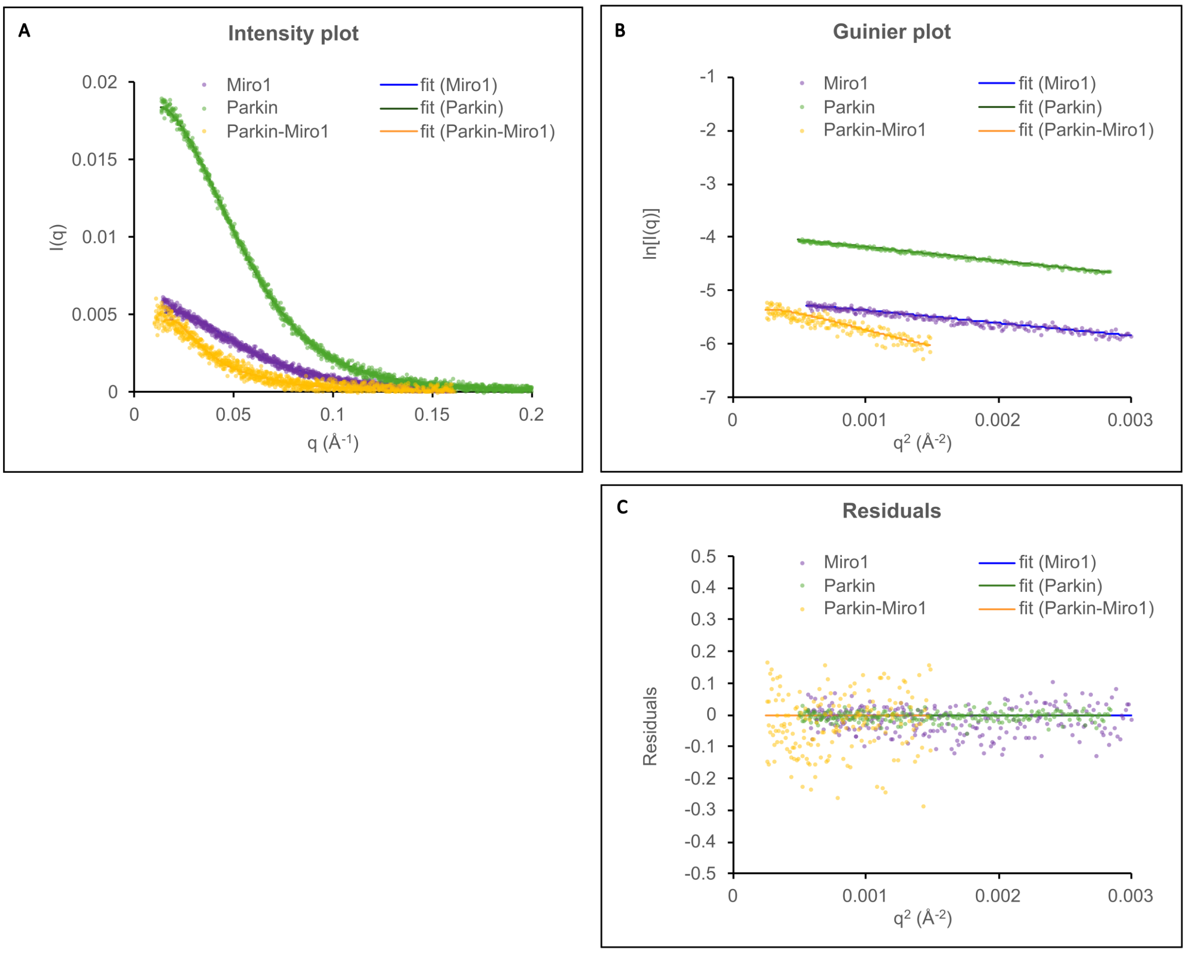


**Supplementary Figure 3: Size exclusion chromatography and small angle scattering (SEC-SAXS) analysis of crosslinked Parkin-Miro1 complex in comparison with Parkin and Miro1 alone.** A: Experimental intensity plots of Parkin (green), Miro1 (purple) and crosslinked Parkin-Miro1 complex (orange). B: Guinier plot of Parkin, Miro1 and crosslinked Parkin-Miro1 complex with residuals shown in C. Rg values of 28 Å for Parkin, 28.5 Å for Miro1 and 39 Å for the complex were derived from these data.

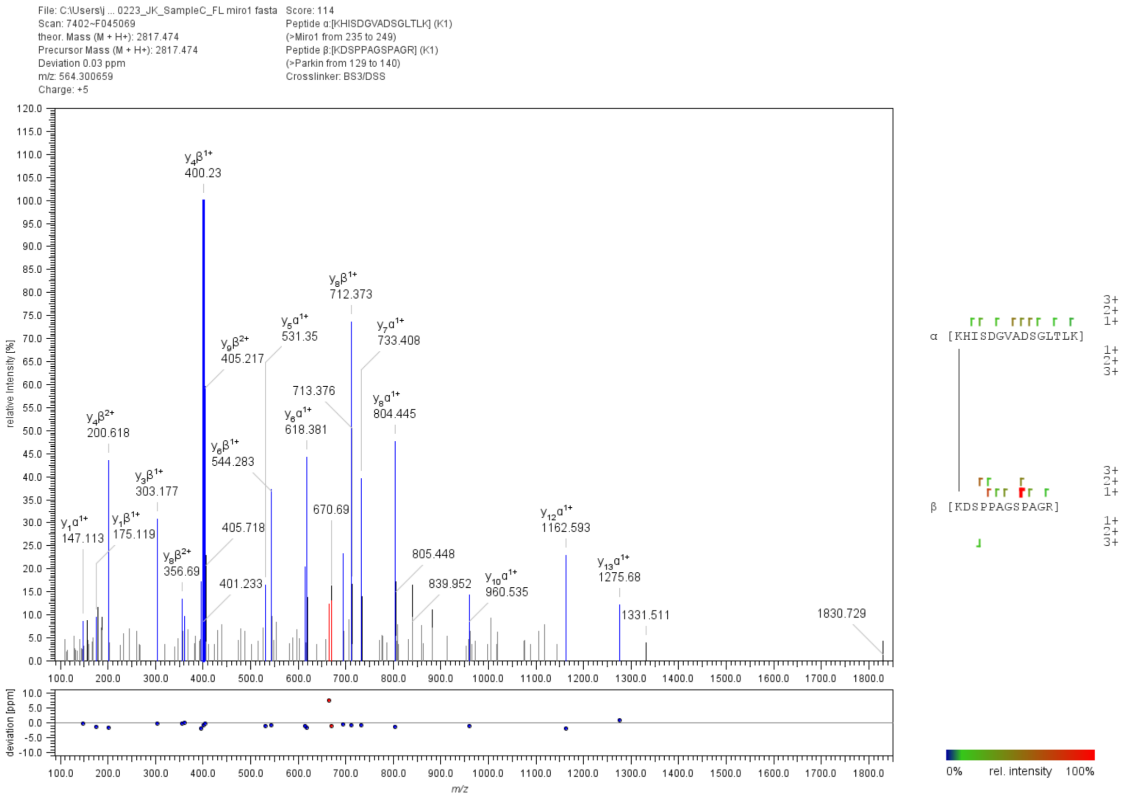


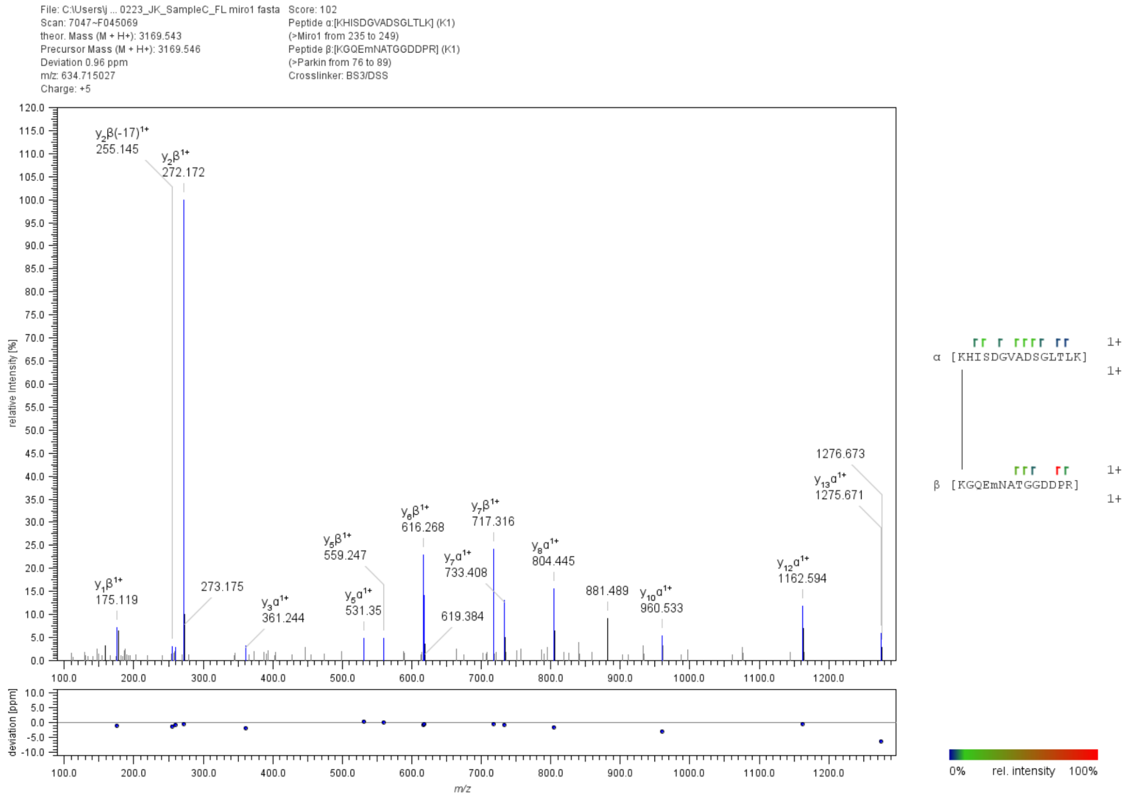


**Supplementary Figure 4: Identification of crosslinked lysines in Parkin-Miro1 complex by mass spectrometry.**

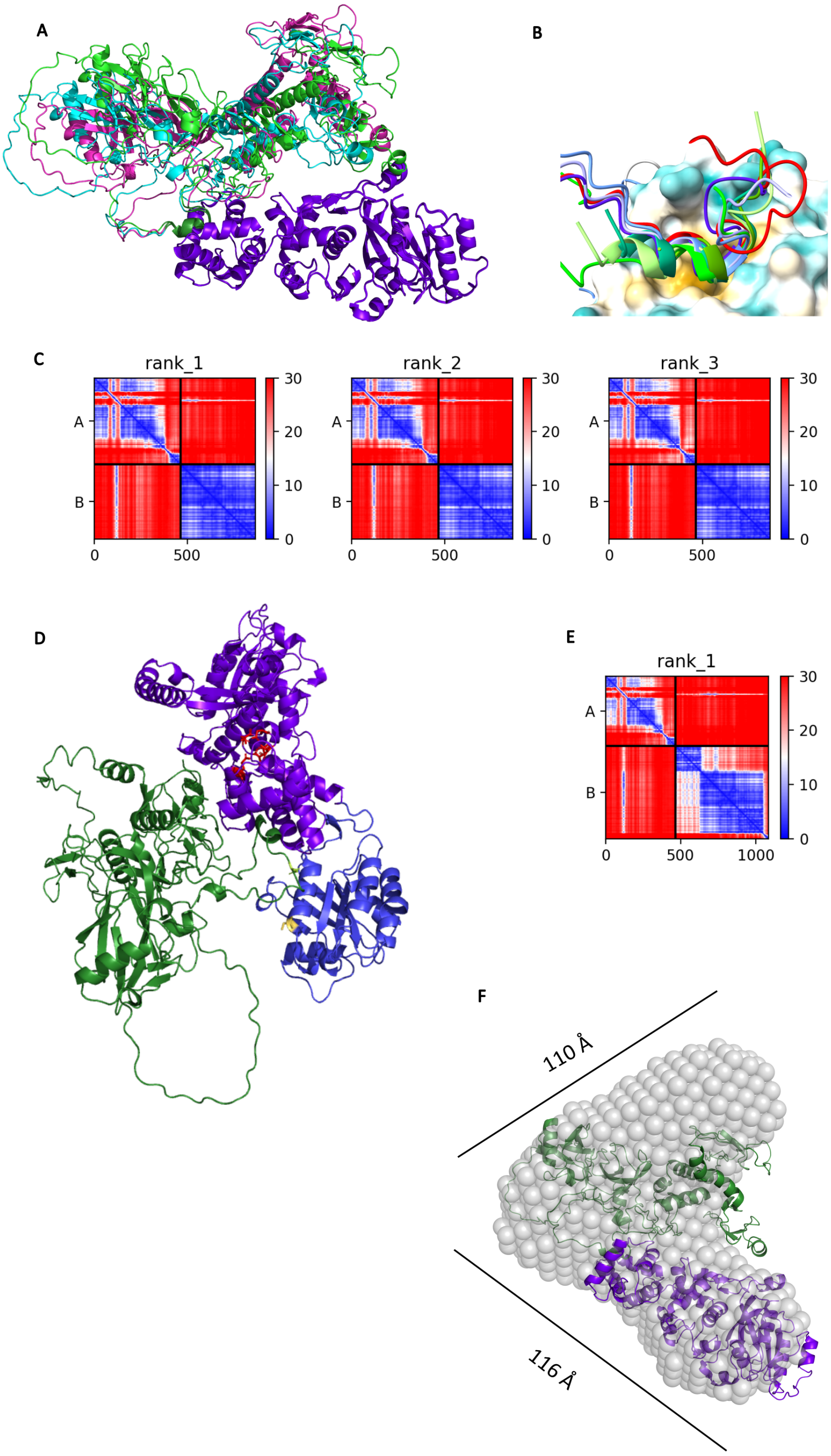


**Supplementary Figure 5: Multiple AlphaFold models predict the same Parkin-Miro1 interacting region with high confidence.** Overlay of three AlphaFold models represented as ribbons, aligned to Miro1 structure (purple). B: Close-up on Parkin-Miro1 interface from A. Miro1 EF1 domain is represented as hydrophobic surface featuring a hydrophobic pocket (yellow). Coloured ribbons represent the Substrate Targeting Region in Parkin (STR). C: PAE plots for three AlphaFold models of Parkin-Miro1 interaction. Parkin STR consistently scores for high confidence score (blue) for its interaction with Miro1. D: AlphaFold prediction of Parkin interaction with full length Miro1. Blue: N-terminal GTPase domain absent in our construct; Yellow stick: S156 reported to be phosphorylated in cells; Red sticks: calcium-binding residues in Miro1. E: Corresponding PAE plot. F: Overlay of the *ab initio* SAXS model with the AlphaFold model.

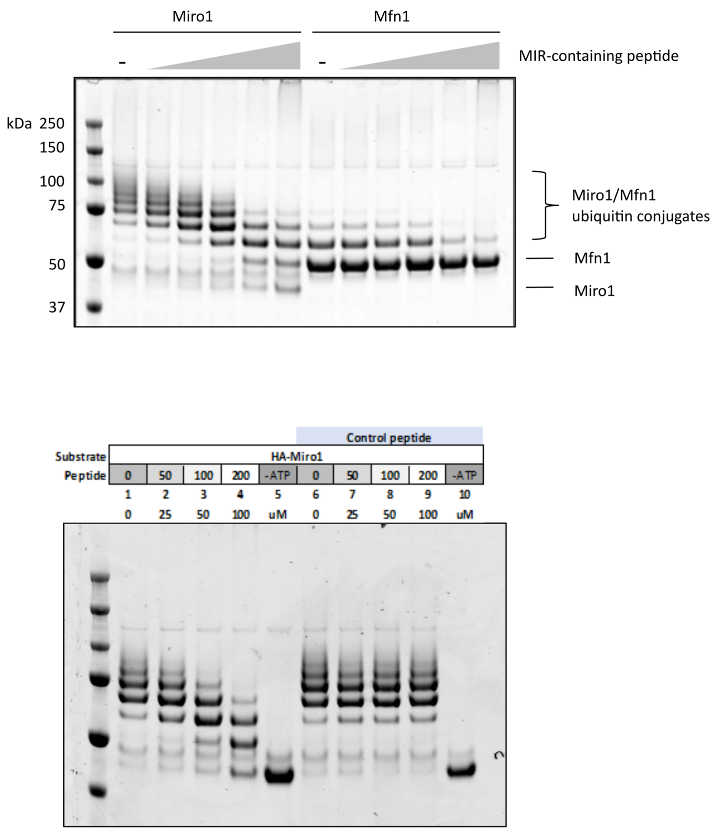


**Supplementary Figure 6: Excess of a synthetic STR-containing peptide decreases Miro1 ubiquitination by Parkin.** A: Miro1 or Mfn1 ubiquitination assays were performed with increasing concentrations of the STR-containing peptide. B: Ubiquitination assays with Miro1 and a control synthetic peptide.
